# A plastid antiporter as a bioindicator of *Thalassiosira pseudonana* resilience

**DOI:** 10.1101/2020.12.09.418384

**Authors:** Jacob J. Valenzuela, Justin Ashworth, Allison Cusick, Raffaela M. Abbriano, E.Virginia Armbrust, Mark Hildebrand, Mónica V. Orellana, Nitin S. Baliga

**Affiliations:** Institute for Systems Biology, Seattle, WA, USA; Scripps Institution of Oceanography, University of California, San Diego, La Jolla, CA; School of Oceanography, University of Washington, Seattle, Washington, USA; Polar Science Center, University of Washington, Seattle, WA, USA; Departments of Biology and Microbiology, University of Washington, Seattle, WA, USA; Molecular and Cellular Biology Program, University of Washington, Seattle, WA, USA; Lawrence Berkeley National Lab, Berkeley, CA, USA

**Keywords:** Diatom, Acidification, Systems Biology, Resilience, Biomarker

## Abstract

Acidification of the ocean due to high atmospheric CO_2_ levels may increase the resilience of diatoms causing dramatic shifts in abiotic and biotic cycles with lasting implications on marine ecosystems. Here, we report a potential bioindicator of a shift in the resilience of a coastal and centric model diatom *Thalassiosira pseudonana* under elevated CO_2_. Specifically, we have discovered, through EGFP-tagging, a plastid membrane localized putative Na^+^(K^+^)/H^+^ antiporter that is significantly upregulated at > 800 ppm CO_2_, with a potentially important role in maintaining pH homeostasis. Notably, transcript abundance of this antiporter gene was relatively low and constant over the diel cycle under contemporary CO_2_ conditions. In future acidified oceanic conditions, dramatic oscillations of >10-fold change between nighttime (high) and daytime (low) in transcript abundances of the antiporter gene were associated with increased resilience of *T. pseudonana*. By analyzing metatranscriptomic data from the Tara Oceans project, we demonstrate that phylogenetically diverse diatoms express homologs of this antiporter across the globe. We propose that the differential between night- and daytime transcript levels of the antiporter could serve as a bioindicator of a shift in the resilience of diatoms in response to high CO_2_ conditions in marine environments.

## INTRODUCTION

Diatoms (Bacillariophyta) represent a diverse group of marine phytoplankton widely distributed across the globe, from coastal habitats to the open oceans (Malviya et al., 2016). They account for 40% of the marine primary productivity (20% globally) (Nelson and Gordon, 1982; Falkowski et al., 1998; Field et al., 1998) and facilitate carbon export to the deep ocean through the biological carbon pump (Falkowski et al., 1998; Smetacek, 1999). Diatoms can form large seasonal blooms when conditions are favorable, generating vast amounts of biomass for marine food webs (Platt et al., 2003). Diatoms have brought stability to key ecosystems (Armbrust, 2009), adjusting to changes in nutrient availability, such as nitrogen (N) (Levitan et al., 2015), phosphorus (P) (Brembu et al., 2017), silicon (Si), iron (Fe) (Marchetti et al., 2009; Smith et al., 2016) and trace elements, as well as physicochemical conditions, including light (Wilhelm et al., 2014; Smith et al., 2016; Lepetit et al., 2017) levels (photosynthetically active radiation; PAR) (Aguirre et al., 2018), temperature (Armbrust, 2009), salinity, pH, and CO_2_ (Falkowski et al., 1998; Raven et al., 2019), over daily, seasonal, and decadal cycles. The robustness of diatoms to combinatorial environmental changes has been investigated through controlled photo-bioreactor studies, demonstrating how the model diatom *Thalassiosira pseudonana* regulates >5,400 genes (>40% of all genes in its genome) to adopt distinct physiologic states matched to relevant environmental conditions (light, dark, nutrient replete and nutrient deplete) (Ashworth et al., 2013). Hundreds of genes encoding sensory, signaling, and regulatory functions execute environment-specific gene regulatory programs to drive transitions of *T. pseudonana* across these physiological states (Ashworth et al., 2013). For example, previous work implicated the photoreceptor AUREO1c in regulation of photosynthesis and sugar metabolism during dawn and exponential growth; while the light-responsive transcription factor bZIP7a/PAS was implicated in regulating other transcription factors. Furthermore, bZIP24b was implicated in regulating nucleosome and chromatin organization at high CO_2_ conditions. Under increasing CO_2_ concentrations, *T. pseudonana* remodels chromatin and transcriptionally downregulates photosynthesis, respiration, and carbon-concentrating mechanisms (CCMs), which are utilized by many diatoms (Badger et al., 1998; Crawfurd et al., 2011) to grow in the CO_2_-limited oceans of today (<10 - 30 μmol CO_2_) (Hopkinson et al., 2011; Reinfelder, 2011).

The reduced need to concentrate carbon at an elevated CO_2_ level alleviates the need for CCMs (Hopkinson et al., 2011; Matsuda et al., 2017), which might allow diatoms to reallocate resources to manage stressful environments. Indeed, at a higher CO_2_ level, *T. pseudonana* was able to withstand incrementally larger doses of ultraviolet radiation (UVR) stress, while transitioning through environment-relevant physiologic states (Valenzuela et al., 2018). Thus, ocean acidification may stabilize diatom populations under NO_3_ limiting conditions and may lead to the expansion of their geographic distribution and ecological niches. However, predicting how diatoms and other phytoplankton will fare in future oceans is complicated by non-linear consequences of combined increases in atmospheric CO_2_, global temperatures, and nutrient availabilities (Feely et al., 2004). For instance, while ocean acidification may improve diatom resilience when they are not Fe limited (Valenzuela et al., 2018), the productivity of diatoms will suffer in environments where Fe bioavailability is predicted to decline with increasing temperature and acidification (Shi et al., 2010). The combined effects of light and CO_2_ on marine productivity can also vary depending on the season, latitude, depth, and the photo-acclimation capabilities of phytoplankton (Jones, 1998), with light intensity setting the upper limit of productivity (Jones, 1998; Platt et al., 2003). Hence, understanding how diatom resilience and productivity will change in response to shifts in the complex interplay of biotic and abiotic factors due to ocean acidification is essential to predict species succession or niche expansions in seasonally acidified environments, such as upwelling systems and in nutrient-limited zones (Bruland et al., 2001; Cohen et al., 2017; Godhe and Rynearson, 2017; Valenzuela et al., 2018).

In order to investigate how varying levels of light will affect diatoms in high and low carbon conditions, we have generated and analyzed the transcriptome responses of *T. pseudonana* to combinatorial changes in light intensity (75 μM photons.m^-2^.s^-1^; hereafter low light or “LL” and 300 μM photons.m^-2^.s^-1^; hereafter high light or “HL”) and CO_2_ level (400 ppm; hereafter low carbon or “LC” and 800 ppm; hereafter high carbon or “HC”). In addition to recapitulating the large-scale physiological state-transitions of *T. pseudonana*, the transcriptome analysis discovered the transcriptional modulation of putative antiporter might play an important role in pH homeostasis during the transition between day and night under elevated CO_2_. In particular, we have discovered a dramatic switch in the dynamical day/night expression patterns of a putative Na^+^(K^+^)/H^+^ antiporter, which indicates a shift in diatom response to high CO_2_ conditions regardless of changes in other environmental conditions (e.g., PAR, UVR, nutrient limitation). Through GFP-tagging we determined that the putative antiporter is localized to the plastid membrane where it might function to help maintain an optimal pH for the activity of enzymes related to carbon fixation, photosynthesis, transport, and other metabolic processes (Launay et al., 2020). This is consistent with the expectation that seawater acidification will alter electrochemical gradients that govern pH gradients across membranes (Taylor et al., 2012). While diatoms typically maintain a neutral pH in the cytoplasm (Taylor et al., 2012; Goldman et al., 2017), the pH within the plastid is higher (pH ∼8) during the light period, and decreases during the dark (pH ∼7) (Launay et al., 2020). This pH homeostasis is essential for plastid function, and therefore, the expression dynamics of the antiporter captures the non-linear consequences of combined changes in multiple environmental factors that influence the overall physiological state of diatoms, making the antiporter a useful composite bioindicator of diatom resilience. By data-mining metatranscriptome sequences from the Tara Oceans project (Carradec et al., 2018; Villar et al., 2018), we determined that phylogenetically diverse diatoms transcribe homologs of this putative antiporter, making it a candidate bioindicator. Importantly, we postulate that it is the differential between night- and daytime transcript expression that will ultimately reveal the shift in diatom resilience in response to elevated CO_2_ associated ocean acidification.

## METHODS

### Experimental design

*Thalassiosira pseudonana* (CCMP 1335) was grown at two CO_2_ levels (400 and 800 ppm) and two light levels, undersaturated (75 μM photons.m^-2^.s^-1^) and saturated (300 μM photons.m^-2^.s^-1^). The cells were pre-acclimated to the respective light conditions before being grown in bottle reactors. RNA samples were collected during the exponential and late-exponential phases of growth. Each experiment was conducted in triplicate, with three serial replicates per combination of CO_2_ and light levels. In total, 24 microarrays were collected and analyzed (8 conditions x 3 replicates each = 24 microarrays).

### Batch culture growth, monitoring, sampling, and analysis

*T. pseudonana* were grown in triplicate batch cultures in 2 L glass bottles containing enriched artificial seawater (ESAW) media with a reduced nitrate concentration of ∼ 100 μM. All cultures were grown under a continuous light regime. During all growth experiments, pH was monitored twice daily (∼ 12 hours apart) using a pH probe calibrated spectrophotometrically, and cell counts were quantified using a hemocytometer. The cultures were equilibrated to 400 ppm or elevated CO_2_ (800 ppm) by bubbling and stirring (50 rpm). The two CO_2_ levels were generated by first stripping the water from the laboratory air with DU-CAL (Drierite company, Xenia, OH, USA) and CO_2_ with Sodasorb (Divers Supply Inc., Gretna, LA, USA) and then using mass-flow controllers (model GFC-17, Aalborg, Orangeburg, NY, USA) to mix with 99.99% pure CO_2_ at exact ratios (Praxair, Danbury, CT, USA). The CO_2_ concentration of the gas was measured with a CO_2_ analyzer (model S151, Qubit Systems, Kingston, Ontario, Canada). The gas flow on each bottle was controlled with a flow meter (model 6A0101BV-AB, Dakota Instruments Inc., Orangeburn, NY, USA). The CO_2_-air mixture was filtered through a 0.2 μm Millipore filter before flowing into the culturing 2 L bottles to maintain axenic cultures. Total dissolved inorganic carbon (DIC) was measured using an Apollo SciTech (DIC analyzer) Model AS-C3 and Li-COR LI-7000 CO_2_/H_2_O analyzer from filtered (0.2 μm) samples. The flasks were inoculated with 5 × 10^4^ cells/mL of acclimated axenic *T. pseudonana* cells. Photosynthetic efficiency (maximal PSII quantum yield, *F*_v_/*F*_m_) was obtained from the maximal fluorescence (*F*_m_) and variable fluorescence (*F*_v_) using the Phyto-PAM (Pulse Amplitude Modulated) Phytoplankton Analyzer (Walz). Variable fluorescence was calculated from *F*_m_ to *F*_o_, where *F*_o_ is the fluorescence yield when cells are dark acclimated. Nutrients (NO_3_, PO_4_, and Si) were sampled twice daily and measured at the Chemical Services Lab (University of Washington).

### RNA extraction and expression microarrays

RNA extraction and expression microarrays were performed according to Ashworth et al. 2013. Total RNA was harvested from a total of 3 x 10^7^ cells from each reactor during a) exponential and b) late-exponential growth, using the mirVANA kit from Invitrogen. A total of 24 RNA samples were collected, based on the following experimental matrix: {400 ppm, 800 ppm} x {unsaturated light, saturated light} x {exponential, late-exponential} x {3 parallel replicates} x {3 serial replicates}. RNA samples were amplified and labeled using the Agilent Quick Amp56 Labeling Kit. The resulting Cy3-labeled cRNA was hybridized along with a uniform Cy5-labeled reference sample (as an internal standard) to custom gene-specific 8 x 60k oligonucleotide Agilent arrays (Agilent design ID: 037886; GEO platform ID: GPL18682). Arrays were scanned using an Agilent two-color array scanner.

### Microarray data processing and gene expression analysis

Two-color microarray data were processed and normalized using the limma package in R (Ritchie et al., 2015). Relative log_2_ expression ratios were calculated in relation to the Cy5-labeled universal standard reference RNA sample. The effects of three dichotomous experimental factors (CO_2_ level, light level, and growth phase) on the expression changes of all genes during the experiments were assessed using a three-way ANOVA in R, with multiple hypothesis correction for significance assessment using the Benjamini-Hochberg methods. Functional enrichment analysis was performed using the g:Profiler toolset (Raudvere et al., 2019). These microarray expression datasets are available in the GEO database under the accession number(s): GSE57737 Series C (n = 24). RNA-seq analysis was performed as described in Valenzuela et al., 2018. In that study, *T. pseudonana* was cultured in photobioreactors under 12:12-hour (light:dark) cycles with nitrate limitation at HC (1,000 ppm CO_2_) and LC (300 ppm CO_2_) to force the transition across 4 physiological states (light, dark, early and late-phase exponential phase). Each growth cycle represented a “stage” and at the end of the first stage an aliquot was transferred to fresh growth medium to initiate the next stage. Starting in stage 2, at midday, 0.5 mW/cm^2^ UVR was applied for 1 hr and incrementally increased by +0.5 mW/cm^2^ in subsequent stages. During each stage, transcriptomes from both HC and LC conditions across the 4 principal states were analyzed for signatures of resilience. These RNA datasets are publicly available from the National Center for Biotechnology (NCBI) Sequence Read Archive (SRA), accession code PRJNA386016.

### Vector construction and *T. pseudonana* transformation

The expression vector used for protein localization was constructed as described in Shrestha and Hildebrand, 2015, using MultiSite Gateway Technology (Life Technologies) (Shrestha and Hildebrand, 2015). The 262258 CDS was amplified with PCR primers flanked with attB sites to insert the fragment into the pDONR221 entry vector. The entry clone facilitated the insertion of the 262258 fragment upstream and in-frame with enhanced green fluorescent protein (EGFP) in the final destination vector pMHL_079, for constitutive transcription under the control of a fucoxanthin chlorophyll a/c binding protein promoter (FCPp). A list of primers used is listed in **Supplementary Table 1**. The FCPp-262258-EGFP-FCPt expression vector was co-transformed with pMHL_009 expressing the nat1 gene under the control of the acetyl-coenzyme A carboxylase promoter, which confers resistance to the antibiotic nourseothricin (NAT). The protocol for *T. pseudonana* transformation was performed as described in Shrestha and Hildebrand, 2015 (Shrestha and Hildebrand, 2015). Following transformation, 250 mL of culture was added to 50 ml of ASW medium containing NAT at a final concentration of 100 ug/mL. The liquid culture was grown to exponential phase (5 x 10^5^ cells/mL) and the top 5% of GFP-expressing cells (10,000 cells total) were sorted by FACS into fresh ASW medium using the BD Influx Cell Sorter. Exponential-phase cells from the sorted culture were used for fluorescence microscopy.

### Confocal fluorescence microscopy

*T. pseudonana* cells expressing the GFP-tagged transmembrane antiporter protein (262258) were imaged with a Zeiss Axio Observer Z1 inverted microscope equipped with an ApoTome and a Zeiss AxioCam MRm camera (Carl Zeiss Microimaging, Inc., Thornwood, NY, USA). The Zeiss #16 filter set was used to image chlorophyll autofluorescence (Excitation BP 485/20 nm, Dichromatic mirror FT 510 nm, Emission LP 515 nm) and the Zeiss #38HE filter set was used to image GFP fluorescence (Excitation BP 470/40 nm, Dichromatic mirror FT 495 nm, Emission BP 525/50 nm). Non-fluorescent images were taken using differential interference contrast (DIC). Z-stack images through the cells were acquired with 63x/1.4 oil immersion Plan-Apochromat objective and a 1.6x optovar module. Images were processed using Axiovision 4.7.2 software. The average composition of 3 Z-stack images was done using imageJ software (Schneider et al., 2012).

### Diatom antiporter sequences in the environment

The Ocean Gene Atlas web server (http://tara-oceans.mio.osupytheas.fr/ocean-gene-atlas/) (Villar et al., 2018) was used to query the Marine Atlas of Tara Oceans Unigenes version 1 Metatranscriptomes (MATOUv1+T) (Carradec et al., 2018) against the amino acid sequence of the putative antiporter (262258). We removed any hits that did not align to the taxonomic phylum Bacillariophyta, and had a bitscore below 300 (largest non-zero e-value 7.69E-107) to ensure sequences were diatom specific with high confidence.

## RESULTS

In order to dissect the combinatorial effects of light and CO_2_, we cultured *T. pseudonana* in four conditions representing combinations of low (75 μM photons.m^-2^.s^-1^, LL) and high (300 μM photons.m^-2^.s^-1^, HL) light levels, and current (400 ppm, LC) and projected (800 ppm, HC) CO_2_ concentrations. We observed similar growth rates across all combinations of light and CO_2_ conditions (HL:HC, HL:LC, LL:HC, and LL:LC; **Fig. 1A**). Cultures in HC achieved a significantly higher carrying capacity (p-value = 0.0027, t-test), as expected (Valenzuela et al., 2018). Nutrient (N, P, Si) uptake rates were also similar across all four treatments, with complete utilization of phosphate within 24 hours (**Supplementary Fig. 1**). Even after depletion of phosphate in the growth medium, the continued growth of diatoms is likely attributable to intracellular polyphosphate reserves (Dyhrman et al., 2012; Valenzuela et al., 2012, 2013) (**Supplementary Fig. 1B**). The complete utilization of all macronutrients by ∼60 hours (i.e., when RNA was sampled for transcriptomic analysis) coincided with a decline in growth rate, a decrease in *F*_v_/*F*_m_ (i.e., maximal fluorescence [*F*_m_] and variable fluorescence [*F*_v_]), and transition of all cultures into late-exponential phase. Consistent with a previous report (Li et al., 2014), *F*_v_/*F*_m_ declined faster during growth in HL regardless of CO_2_ level (**Fig. 1B**). DIC decreased proportionally with an increase in biomass, and DIC levels were restored during transition to stationary phase when nutrients were depleted, and PE decreased (**Fig. 1C**).

**Figure 1.**
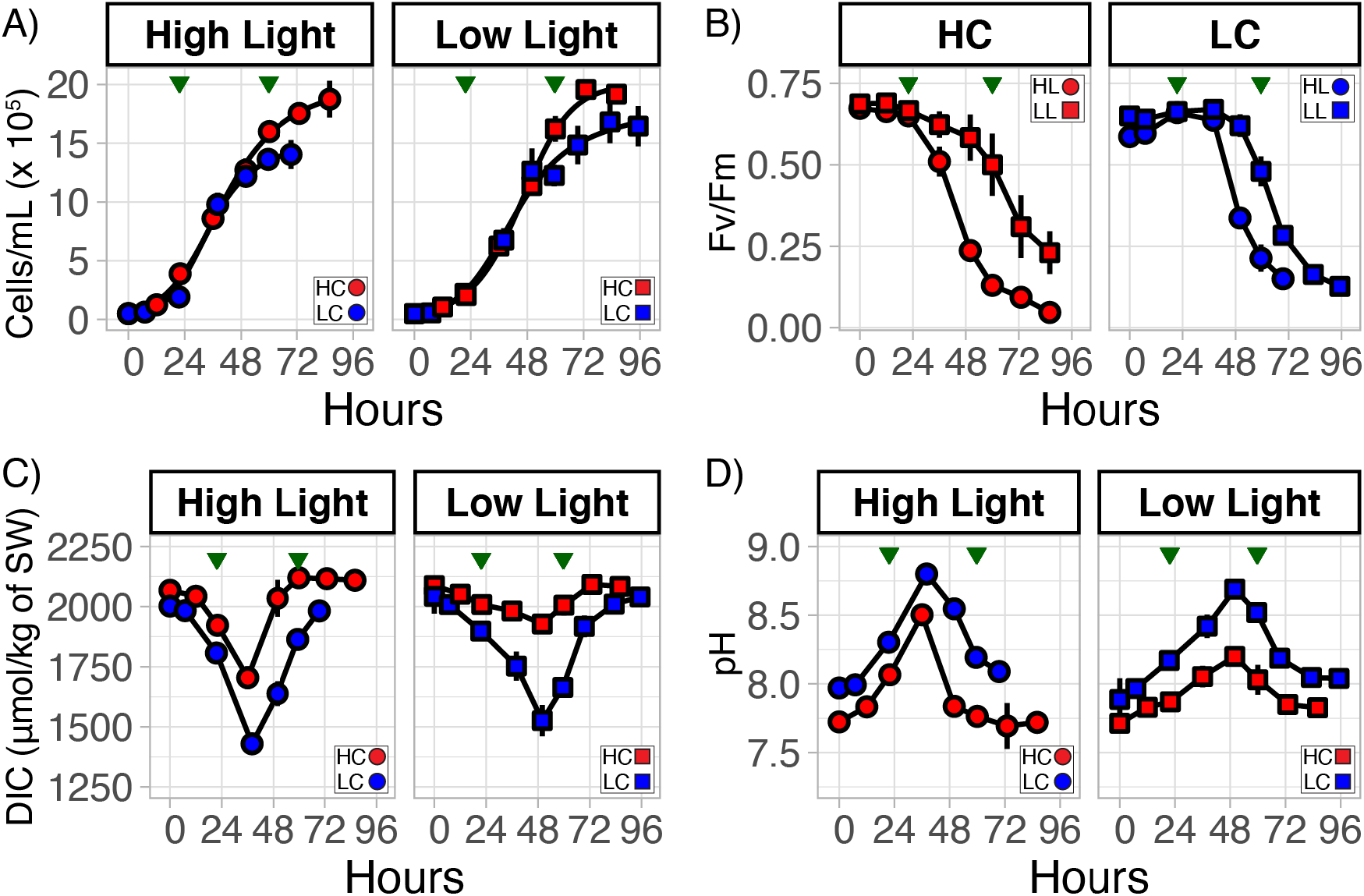
Growth dynamics of *T. pseudonana* with combinatorial changes in light intensity and CO_2_ level. (**A**) *T. pseudonana* growth at low light (LL) and high light (HL) at elevated (800 ppm, HC, red) and low (400 ppm, LC, blue) CO_2_ conditions. **(B)** Dynamics of growth-associated longitudinal changes in photosynthetic efficiency (*F*_*v*_/*F*_*m*_), (**C**) dissolved inorganic carbon (DIC), and (**D**) pH at HL:HC, HL:LC, LL:HC, and LL:LC conditions. Error bars correspond to the standard deviation (n = 3).

Uninterrupted photosynthesis due to culturing in continuous light conditions did not allow for external pH recovery in the media, which typically occurs during nighttime (i.e., no photosynthesis) (Ashworth et al., 2013; Valenzuela et al., 2018). As a consequence, pH continued to increase in all cultures through late-exponential phase. In general, the rate of increase and level of pH was higher in HL conditions irrespective of CO_2_ levels. HC cultures maintained relatively lower pH throughout the experiment. The highest pH (∼8.75) was observed in the HL:LC condition; by contrast, LL:HC reached a maximum pH of ∼8.2. Notably, the direction of change in pH flipped when cultures transitioned into late-exponential phase, which happened earlier in HL (∼36 hours), relative to LL (∼48 hours) grown cultures, and coincided with a decrease in PE. As expected, changes in DIC and pH of the culture media were inversely correlated (**Fig. 1C and D**). In other words, growth characteristics of diatoms in these experiments resembled bloom dynamics, where drawdown of nutrients can be attributed to growth of diatoms, simultaneously causing large fluctuations in daily pH (>1 pH unit) over the light and dark periods (Wallace et al., 2014; Raven et al., 2019).

We performed whole transcriptome analysis on cells harvested during the mid- (∼24 hrs) and late-exponential (∼60 hrs) phases of growth to investigate molecular and systems level changes associated with the response of diatoms to combinatorial changes in light and CO_2_. Using three-way analysis of variation (ANOVA) we identified over 1,700 genes with greater than a two-fold change in transcript abundance (FDR <= 0.001; **Table 1, Supplementary Data File 1**). This analysis also allowed attribution of differential regulation of genes due to changes in each of the three variables in this experiment (growth phase, light intensity and CO_2_ level). As expected, transition from mid- to late-exponential growth phase accounted for the majority of transcriptional changes. The 884 genes that were upregulated in mid-exponential phase to support growth and biomass production were subsequently downregulated in late-exponential phase when nutrients were depleted. Functions enriched within these genes included photosynthesis (e.g., light-harvesting complexes and photosystems I and II reaction center proteins) and carbon fixation (e.g., RuBisCO large subunit [bd2088], glyceraldehyde 3-phosphate dehydrogenase [31383], and ribose-5-phosphate isomerase [32252]) (**Supplementary Fig. 2A and B**). By contrast, 816 genes were downregulated at mid-exponential phase (upregulated at late-exponential phase) and associated with functions related to ribosome metabolism and nutrient transport (**Supplementary Fig. 2C and D**). In particular, the upregulation of nitrate, phosphate, and silicic acid transporters (269274, 27414, 261414, and 268895) was consistent with the depletion of these micronutrients in stationary phase (Mock et al., 2008; Valenzuela et al., 2012; Ashworth et al., 2013; Lomana et al., 2015). The coordinated upregulation of multiple RNA helicases, a ubiquitin-conjugating enzyme (36045), and an endonuclease (bd410) suggests that *T. pseudonana* may utilize these genes to recycle N and P from nucleotides (Bourgeois et al., 2016; Brembu et al., 2017; Alipanah et al., 2018) during nutrient starvation-induced growth arrest. In sum, transition of cells into late-exponential phase was associated with downregulation of energy production and upregulation of RNA metabolism and nutrient transport, suggesting a shift towards energy conservation and nutrient scavenging.

**Table 1.** Relative transcriptome changes in different light and CO_2_ levels. Significant transcriptome changes were determined with three-way ANOVA (CO_2_ level, light level, and growth phase). Number of gene models with transcriptional changes that were significantly associated with CO_2_ level, light level, and growth phase. Bold numbers in parentheses indicate the number of these significant changes that were greater than two-fold. ^1^Benjamini-Hochberg was applied to all ANOVA *p*-values with *n* = 15,059.

| **Light Intensity Experiment (24 microarrays):**<br>Diatoms pre-acclimated to either 300 or 75 $\mu\text{M}$ photons$\cdot\text{m}^{-1}\cdot\text{s}^{-1}$ prior to experiments; light intensity modulated using neutral density filters | **Number of significant transcriptional changes** | | |
| --- | --- | --- | --- |
| | FDR<sup>1</sup> ≤ 0.01 | FDR<sup>1</sup> ≤ 0.001 | FDR<sup>1</sup> ≤ 0.0001 |
| Growth Phase (Exponential vs. Stationary) | 6,335 (**1,833**) | 4,438 (**1,700**) | 3,004 (**1,496**) |
| Light Intensity (300 vs. 75 $\mu\text{M}$ photons$\cdot\text{m}^{-2}\cdot\text{s}^{-1}$) | 96 (**27**) | 27 (**16**) | 12 (**10**) |
| CO<sub>2</sub> (800 vs. 400 ppm) | 2,024 (**145**) | 240 (**36**) | 10 (**9**) |

Interestingly, only 16 genes had a greater than a two-fold differential regulation (FDR <= 0.001) as a direct consequence of the change in light intensity (6 upregulated in HL and 10 in LL). Similarly, differential regulation of just 36 genes could be attributed to CO_2_ level change, with 16 genes upregulated in HC (**Fig. 2A**) and 20 genes upregulated in LC (**Table 1 and Supplementary Data File 1**). While there was no significant enrichment of known functions among the differentially regulated genes, the analysis did recapitulate high CO_2_-responsive downregulation of 3 genes [6528, 10360 (a putative transcriptional regulator: Tp_bZIP24a) and 233 (a putative carbonic hydrase)], that were previously identified as part of a CCM and photorespiration sub-cluster (Ashworth et al., 2015; Hennon et al., 2015) (**Supplementary Table 2 and Supplementary Data File 1**). Among the five most upregulated genes at high CO_2_ level, three had unknown functions and two had putative transporter functional annotations (**Supplementary Table 2**). We investigated the robustness of CO_2_-responsive upregulation of the five genes by analyzing an independent RNA-seq dataset from a study designed to investigate the resilience of *T. pseudonana* at two CO_2_ levels (300 ppm and 1,000 ppm) over the diel cycle (Valenzuela et al., 2018). In brief, *T. pseudonana* cultures were grown under a 12:12-hour light:dark cycle in nitrogen-limited growth medium, and an aliquot from late-exponential phase was inoculated into fresh medium –each growth cycle representing a “stage”. In stage 2, the cultures received one hour of an ecologically relevant dose of UVR (0.5 mW/cm^2^) at midday; and incrementally higher doses (+0.5 mW/cm^2^) in each subsequent stage, until cultures collapsed. This study demonstrated increased resilience of *T. pseudonana* in HC conditions, which manifested in their ability to recover from the increased number of exposures at a higher dose of UVR through stage 3 (presumably by reallocating energy savings from downregulation of CCMs (Hopkinson et al., 2011)), unlike LC cultures, which collapsed in stage 3. Analysis of transcript level changes and absolute expression (FPKMs: Fragments Per Kilobase of transcript per Million) of the five genes in RNA-seq data from all three stages of this experiment demonstrated that only the putative antiporter (262258) was significantly differentially expressed with a relatively high absolute transcript abundance. The other four genes had relatively low transcript abundance (**Supplementary Fig. 3**) and only gene 5647 was significantly upregulated in HC conditions. The transcript abundance of the putative antiporter was higher at 1,000 ppm CO_2_ by up to 52-fold (mean = 129.3, max = 251.4 FPKMs) relative to its transcript level in 300 ppm CO_2_ (mean = 2.49, max = 8.29 FPKMs) (**Fig. 2B, Supplementary Data File 2**). The difference in the transcriptional patterns of the other four transcripts across the two studies suggests that the regulation of these genes may be sensitive to constant light vs diel conditions. Thus, the comparative analysis of *T. pseudonana* transcriptome responses to high and low CO_2_ in different contexts (i.e., with constant light at two different intensities [using microarrays, this study] and over diel conditions with intermittent UVR stress [RNA-seq] (Valenzuela et al., 2018)), demonstrated that diel changes in abundance of the antiporter transcript might serve as a robust predictor of a shift in the resilience of diatoms in response to elevated CO_2_ in the natural environment, irrespective of changes in other factors.

**Figure 2.**
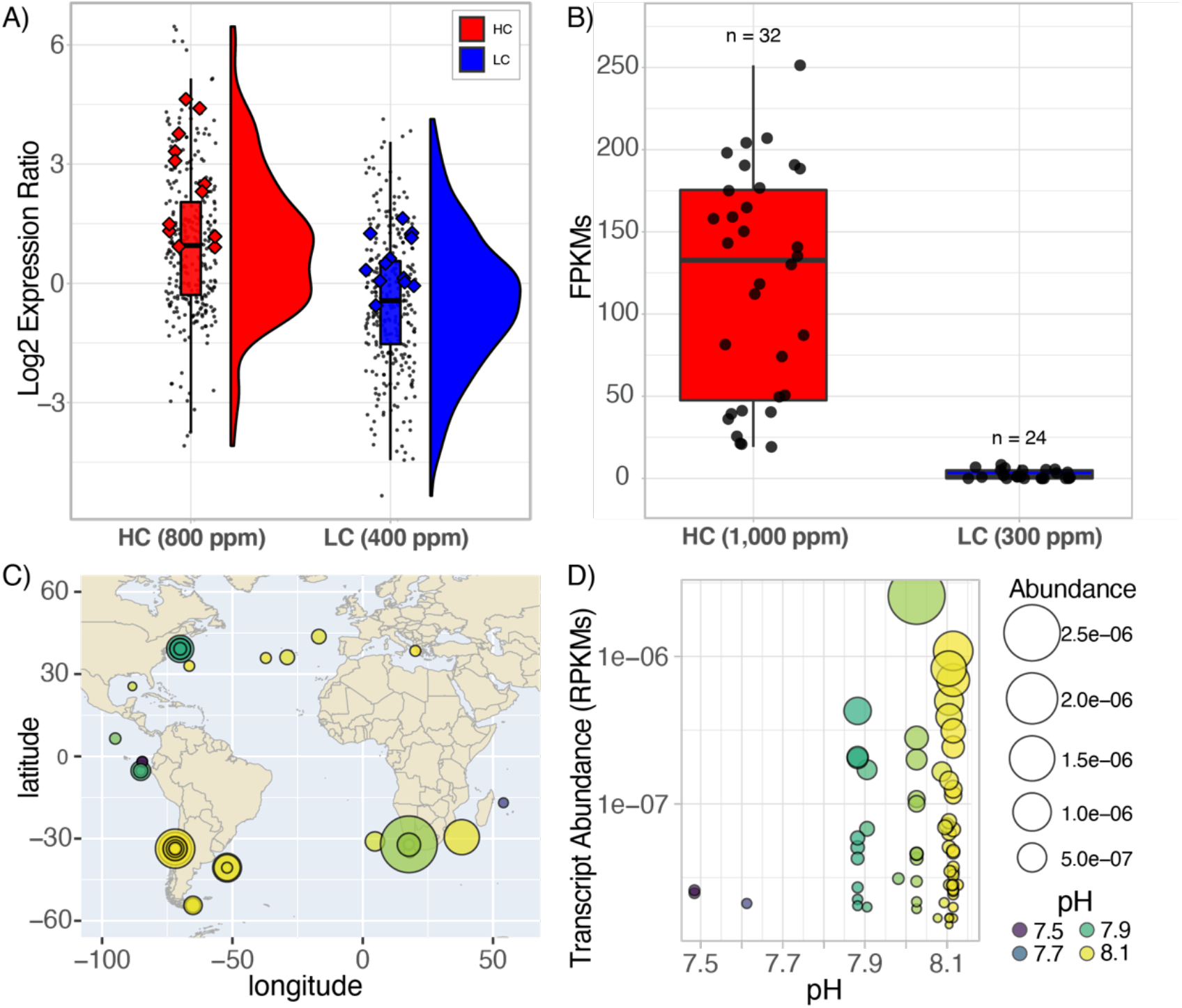
Elevated expression of the putative antiporter (262258) at HC conditions from two different studies and the relative abundance of antiporter homologs across the globe. (**A**) Rain-plots and histograms representing the distribution of expression of 16 upregulated CO_2_-responsive genes. Each dot represents the replicates at HL and LL for both exponential and late-exponential phase at HC (red) and LC (blue) conditions for each gene. Enlarged diamond markers indicate expression values for the putative antiporter (262258). (**B**) Absolute expression values (FPKMs) for the putative antiporter over multiple diel and growth cycles (stages) of a stress test demonstrate upregulation at HC (red) vs. LC (blue) conditions (p-value= 0.00005; Supplementary File 2, Valenzuela et al. 2018). (**C**) Permeases sequences are widely distributed across the globe, in particularly at coastal regions. The area of each circle reflects the abundance values for the sequence at that site (larger area = higher abundance). The color scheme reflects the pH of the environment the sample was collected. (**D**) Transcript abundance of the antiporter homolog sequences from the query of the MATOUv1+T dataset were observed at high and low pH conditions, which was negatively correlated with CO_2_ conditions.

We further investigated whether the putative antiporter and its homologs in other diatoms were also expressed in the natural environment by querying the Marine Atlas of Tara Oceans Unigenes Eukaryotic Metatranscriptomic database (MATOUv1+T) using the Ocean Gene Atlas webserver (Villar et al., 2018). We ascertained diatom-specificity of antiporter homologs by imposing stringent filters to eliminate potential homologs that did not align to the taxonomic phylum Bacillariophyta or had a bitscore below 300 (equated to the largest non-zero e-value 7.69E-107). Altogether 66 diatom-specific antiporter sequences were identified with high confidence, of which 6 homologs were attributed to the genus *Chaetoceros*, the most abundant diatom genus in the world (Rines and Theriot, 2003; Malviya et al., 2016). The diatom antiporter transcripts were discovered in ocean samples across the globe, mostly in coastal regions, and exclusively in the surface water layer (**Fig. 2C**). We inferred relative abundance of each putative antiporter transcript at a specific location as the percentage of all mapped reads in the associated samples (RPKMs: Reads Per Kilobase covered per Million of mapped reads) (Villar et al., 2018). The wide variation in abundance of transcripts in samples of similar pH, including samples from the same location, demonstrated that absolute transcript level of the antiporter “alone” may have no predictive value, vis-à-vis the status of CO_2_-responsiveness of diatoms at a given location (**Fig. 2D**). Therefore, using the same diel dataset from Valenzuela et al. 2018, described above, we investigated whether dynamic changes in the expression level of the antiporter transcript (262258) over the diel cycle was diagnostic of a shift in the resilience of *T. pseudonana* in future oceanic conditions. The antiporter expression level did not change significantly over the diel cycle at 300 ppm CO_2_ and had low absolute expression. Strikingly, at 1,000 ppm CO_2_, the transcript level of the permease oscillated dramatically with low daytime levels (19.2 FPKMs) and high nighttime levels (251.4 FPKMs) (**Fig. 3A**). These findings suggest that the increased resilience of *T. pseudonana* could be inferred from the differential in daytime and nighttime expression levels of the antiporter particularly when functioning at elevated CO_2_.

**Figure 3.**
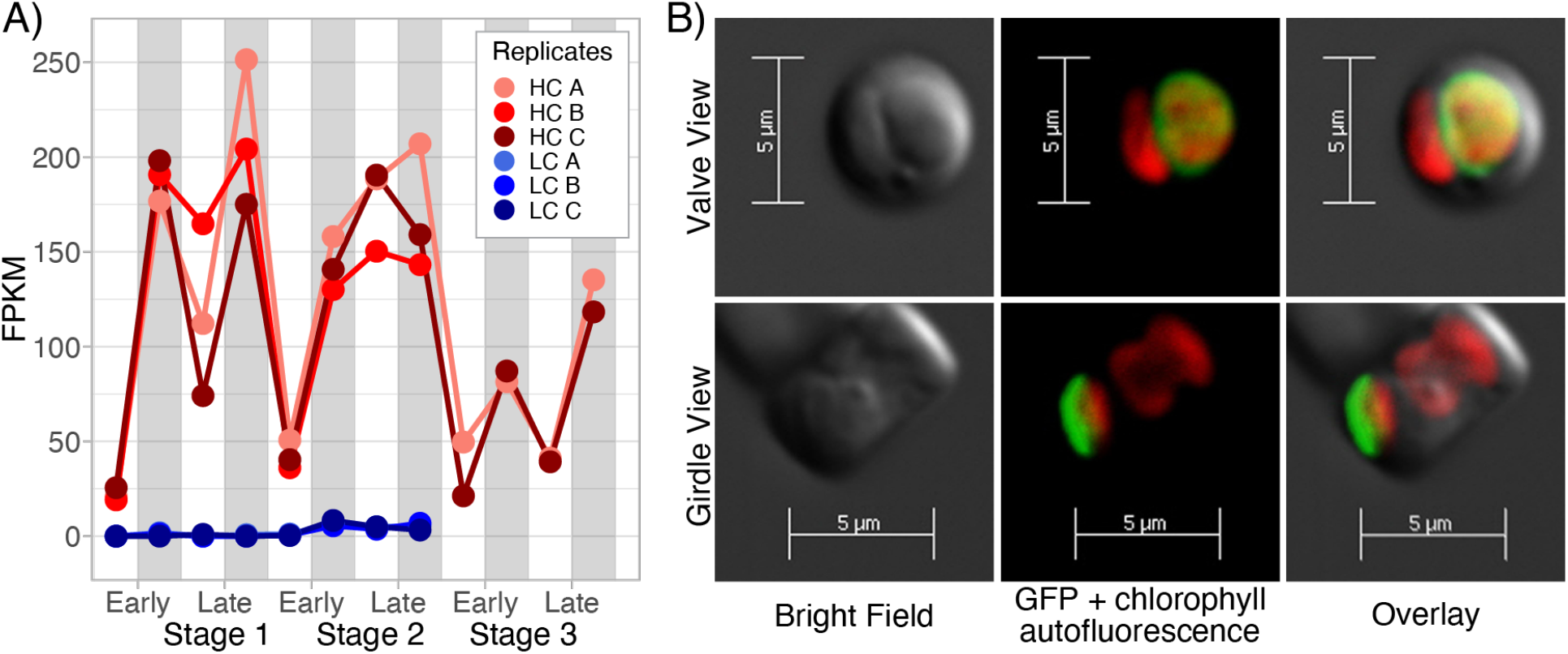
Dynamic diel cycle expression pattern for the putative antiporter and its localization to the plastid. (**A**) Line plots of absolute expression demonstrate the dynamic daytime (white bar; low expression) and nighttime (gray bar; high expression) aligned oscillations in HC (red) but not LC (blue) conditions, with maximal expression in nighttime and late-exponential phase of growth, when nutrients were depleted. (**B**) Bright field and confocal fluorescence microscopy images of two cells of *T. pseudonana* expressing the EGFP-tagged putative antiporter 262258, from the valve and girdle perspectives. Integrated z-stack view of three focal planes shown in green (470/40 nm excitation) that EGFP-tagged antiporter is localized to the membrane surrounding the plastid. The red color is chlorophyll autofluorescence (485/20 nm excitation). Bright-field images were taken using differential interference contrast.

We sought to understand the putative role of the antiporter that may explain the dramatic switch-like shift in its dynamic expression patterns over the diel cycle as a function of increased resilience of diatoms in response to a shift from low to high CO_2_. Secondary structure, organelle localization (TargetP) (Emanuelsson et al., 2000), and functional domain classifications (GO0005215: transporter activity; KOG2639: inorganic ion transport), suggested that the putative antiporter (262258) is a plastid-localized (**Supplementary Table 3**), 12-transmembrane Na^+^/H^+^ antiporter (Wang et al., 2018b, 2018a) (**Supplementary Fig. 4**). In order to confirm its localization to the plastid membrane, we chromosomally integrated an engineered DNA construct for constitutive expression of a C-terminal EGFP-tagged antiporter (see Methods for details) to interrogate its intracellular distribution using confocal fluorescence microscopy (**Fig. 3B**). Analysis of the fluorescence and confocal images confirmed the predicted localization of the transmembrane antiporter to the plastid membrane. Notably, we discovered that the ortholog of the Na^+^/H^+^ antiporter in the model pennate diatom, *Phaeodactylum tricornutum* (Phat3_EG02645, also a plastid localized transmembrane protein with putative inorganic ion transport activity) is also upregulated in the dark with low expression in light (Smith et al., 2016) (**Supplementary Table 3 and Supplementary Fig. 5**). Together, the ∼5-fold upregulation (FDR = 1.1E-04) and dynamic day/night oscillations in HC conditions, localization to the plastid membrane, and putative Na^+^/H^+^ antiporter function suggested that the transmembrane protein may potentially play an important role in maintaining electrochemical gradients across the plastid membrane in the context of diel cycles and elevated CO_2_ levels (Hennon et al., 2015). If so, then the function of the antiporter might become even more essential in an acidified ocean, further supporting its potential utility as a bioindicator of a shift in diatom resilience in the natural environment.

## DISCUSSION

In this study, we observed that >1,700 genes were differentially regulated (FDR > .001) during the growth of *T. pseudonana* in combinations of high light (HL), low light (LL), high carbon (HC), and low carbon (LC), which simulated bloom-like dynamics of rapid growth in nutrient-replete conditions followed by a transition to a more quiescent state in nutrient-deplete stationary phase. The discovery that relatively few genes were differentially expressed exclusively because of changes in individual factors demonstrated the degree of combinatorial regulation of diatom response by light, CO_2_ and nutrient availability. It is noteworthy that the differential expression of the putative antiporter was directly reflective of response to CO_2_, across different intensities of light, UV stress and nutrient availability. We postulate that this is because the gene is potentially an intracellular pH-sensitive Na^+^(K^+^)/H^+^ antiporter with a direct role in maintaining pH homeostasis across the plastid membrane, similar to the chx23 Na^+^(K^+^)/H^+^ antiporter in *Arabidopsis thaliana*, which by itself has been shown to maintain pH homeostasis in the chloroplast stroma (Song et al., 2004). In fact, *A. thaliana* also uses a K^+^/H^+^ antiporter to modify the proton motive force in the thylakoid lumen during the light phase (Kunz et al., 2014; Finazzi et al., 2015).

During dark to light transitions, in many plants as well as diatoms, the internal pH of the plastid can increase from ∼7 to ∼8 (Colman and Rotatore, 1995), which is the optimal pH for the activity of most plastid enzymes, including RuBisCO and Calvin cycle enzymes (Goldman et al., 2017). Given that the cytosolic pH of diatoms in contemporary CO_2_ levels is in the range of 7.0 to 7.5 (Taylor et al., 2012; Höhner et al., 2016; Goldman et al., 2017; Launay et al., 2020), active efflux of H^+^ or counter exchange of ions by means of antiporters may be necessary to achieve a pH of 8 inside the plastid. Studies on *T. weissflogii* have demonstrated that acidification of seawater proportionally decreases intracellular pH causing internal acidosis in diatoms (Goldman et al., 2017). By 2021, increased atmospheric CO_2_ is expected to drive down the pH of seawater from to 7.6 (Caldeira and Wickett, 2005), which may disrupt the intracellular acid-base balance of the plastid and other organelles. The switch from low and non-dynamic day/night expression profile of the antiporter at 300 ppm CO_2_ to dramatic oscillations at 1,000 ppm CO_2_ suggests that in acidified oceans, the role of the putative Na^+^(K^+^)/H^+^ antiporter might become even more important in regulating pH homeostasis during diel transitions. Absolute transcript level of the antiporter at a given location is proportional to the relative abundance of diatoms in that location, and their response to their environmental context (e.g., nutrient availability, CO_2_ level, light condition, etc.). Therefore, absolute transcript level alone is not indicative of the CO_2_-response of the diatom. Instead, we postulate that the differential between nighttime and daytime expression of the antiporter is a better proxy for the overall physiologic response of the diatom to non-linear consequences of changes in multiple factors associated with ocean acidification. Importantly, the switch in expression pattern of the putative antiporter from a non-dynamic to dynamic oscillations may inform when specific diatom species in a given environment are likely to transition to a more resilient state, potentially driving a shift in ecosystem dynamics of a marine habitat.

## ACKNOWLEDGMENTS

We thank the National Science Foundation (Grants MCB-1316206 to M.V.O. and N.S.B.; DBI-1262637, DBI-1565166, MCB-1330912, and MCB-1616955 to N.S.B.) and the National Institutes of Health (Center award 2P50GM076547 to the Institute for Systems Biology). We would like to thank Pamela Troisch and Danielle Yi in the Molecular and Cell core at the Institute for Systems Biology for their help in microarray processing procedures. R.A. was supported by the Department of Energy Office of Science Graduate Fellowship Program (DOE SCGF), made possible in part by the American Recovery and Reinvestment Act of 2009, administered by ORISE-ORAU under contract no. DE-AC05- 06OR23100. All authors would specifically like to thank Mark Hildebrand who unfortunately passed away during this study. M.H touched many lives both personally and scientifically, he was a compassionate leader and gracious colleague.

## AUTHOR CONTRIBUTIONS

J.A., A.C., E.V.A., M.V.O., and N.S.B. designed the experiments, which were performed at E.V.A’s laboratory. J.A., A.C., M.V.O., performed all growth experiments, assays, RNA extraction, and preparation. J.A. and J.J.V. performed microarray and transcriptome analyses. J.J.V. performed meta-transcriptomic and additional RNA-seq analyses. M.H. provided the laboratory and resources for R.M.A. to construct all vectors and perform fluorescence microscopy. J.J.V., J.A., A.C., R.M.A, E.V.A., M.V.O., and N.S.B contributed to the discussion of results. J.J.V., M.V.O, J.A., and N.S.B. wrote the manuscript.

## CONTRIBUTION TO THE FIELD STATEMENT

Diatoms account for 40% of the marine primary productivity and facilitate carbon export to the deep ocean. Ocean acidification due to high atmospheric CO_2_ levels may increase the resilience of diatoms causing dramatic shifts in abiotic and biotic cycles with lasting implications on marine ecosystems. Knowledge of when such shifts are likely to occur has important implications in proactive ecosystem management. However, predicting how diatoms and other phytoplankton will fare in future oceans is complicated by non-linear consequences of combined effects of atmospheric CO_2_, global temperatures, and nutrient availability. Here, we report a putative plastid transmembrane antiporter that could serve as a bioindicator for detecting an ocean acidification-driven shift in the resilience of a key coastal and centric diatom *Thalassiosira pseudonana*. Notably, we have discovered that a shift in the resilience of *T. pseudonana* from current (∼400 ppm CO_2_) to future oceanic conditions (>800 ppm CO_2_) is associated with a transition in expression pattern of the antiporter from constitutive low levels to dynamic oscillations, with >10 fold higher expression at nighttime. Thus, expression dynamics of the antiporter could serve as an early warning sign of marine communities approaching a critical transition, enabling proactive risk mitigation or ecosystem preservation strategies.

