## Supplementary Material for "A plastid antiporter as a bioindicator of *Thalassiosira pseudonana* resilience"

### SUPPLEMENTARY FIGURES AND TABLES

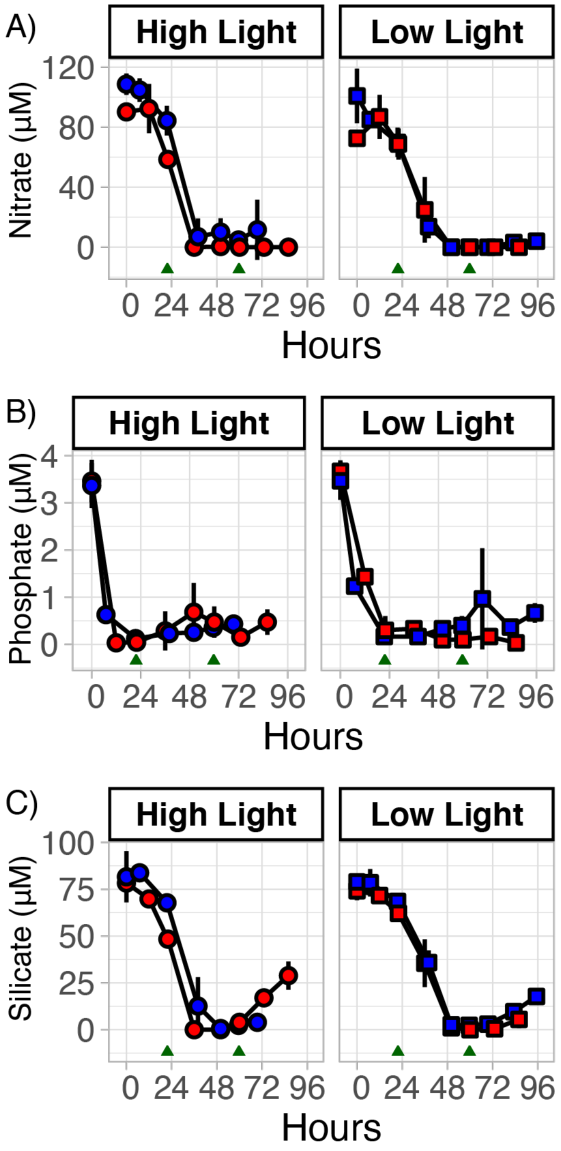

**Supplementary Figure 1. Nutrient drawdown across all conditions during growth with continuous light showed no differences between light or carbon condition.** (**A**) Nitrate, (**B**) phosphate, and (**C**) silicate levels and depletion rates across different light intensities and $CO_2$ levels. Green triangles indicate when cells were harvested for transcriptome profiling (microarray) across the exponential and late-exponential phase of growth.

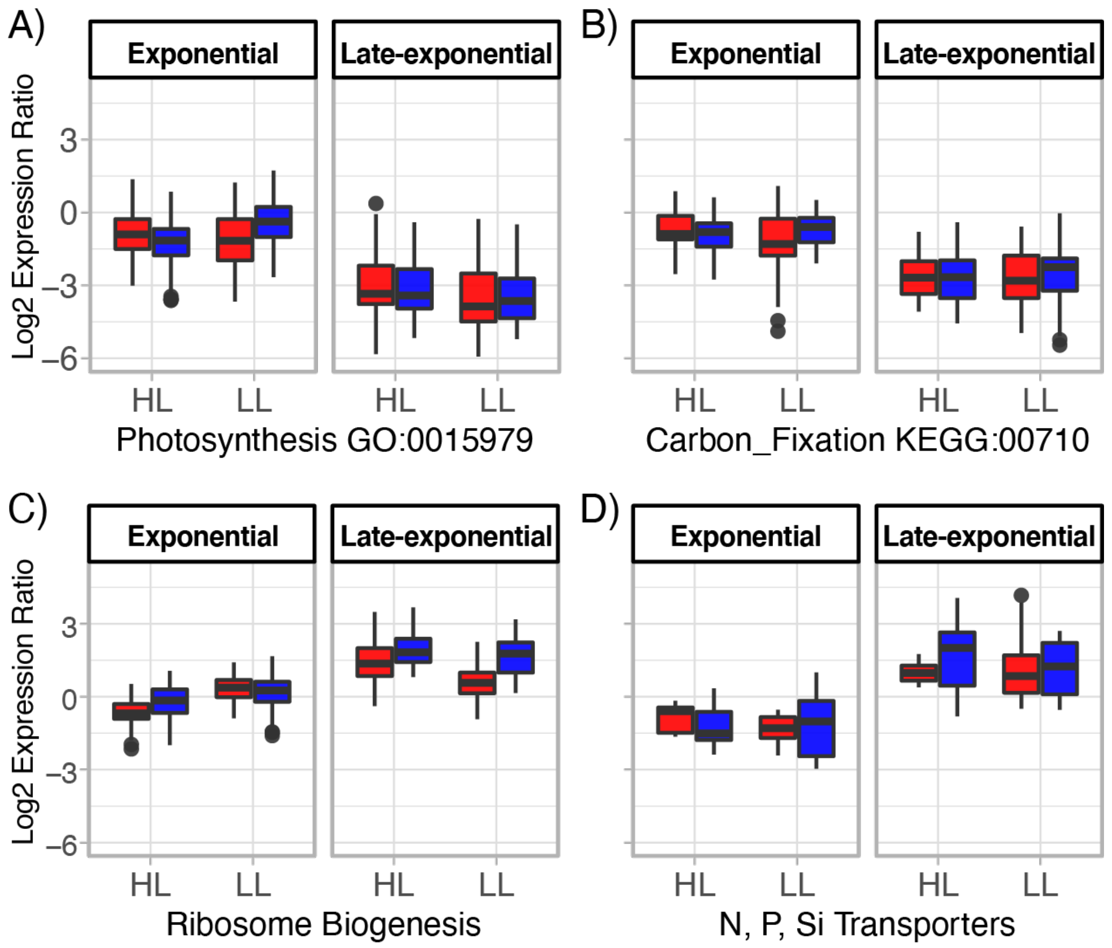

**Supplementary Figure 2. Gene expression dynamics for functional clusters that are significantly differential expressed between exponential and stationary phase.** (**A**) Photosynthetic (*n* = 48) and (**B**) carbon fixation (*n* = 11) genes were expressed at a higher level during exponential phase, but no significant differences were observed across different light intensities or $CO_2$ conditions. (**C**) Ribosome biogenesis (*n* = 32) and (**D**) nutrient transporters (*n* = 4) were upregulated during late-exponential phase, with no significant differences across different light intensities or $CO_2$ conditions.

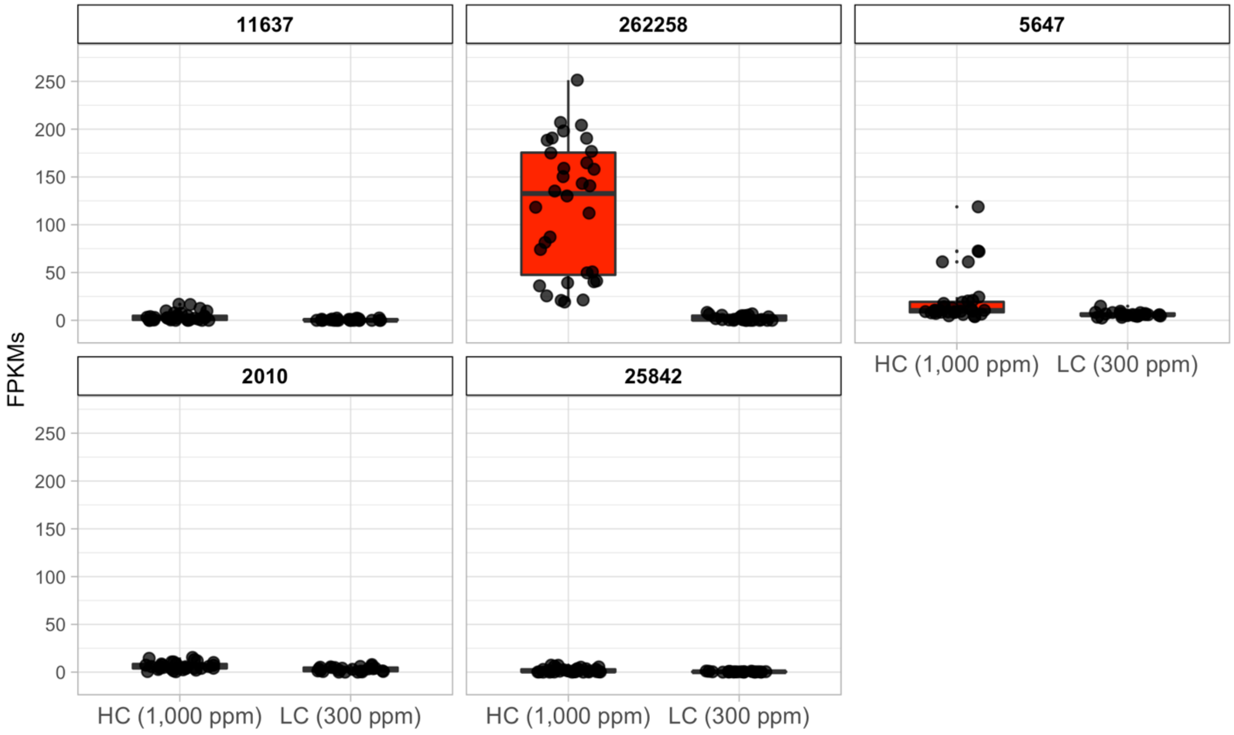

**Supplementary Figure 3. Absolute expression (FPKMs) of the top five significantly differentially upregulated genes (Table 1) using the RNA-seq data of a study designed to investigate the resilience of *T. pseudonana* at two $CO_2$ levels (300 ppm and 1,000 ppm) over the diel cycle.** FPKMs are plotted on the same scale to allow for comparison of transcript levels of all 5 genes relative to each other.

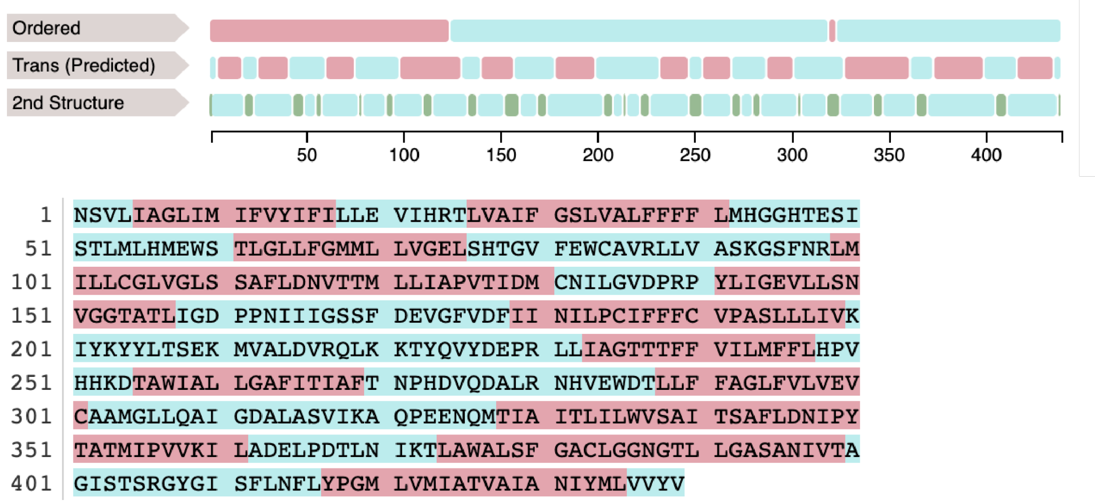

**Supplementary Figure 4. Transmembrane prediction of the putative antiporter using PredMP.** The top annotation is the predicted results of ordered/disordered regions, transmembrane topology (red), and secondary structure. The bottom annotation is the antiporter amino acid sequence with the transmembrane regions highlighted in red.

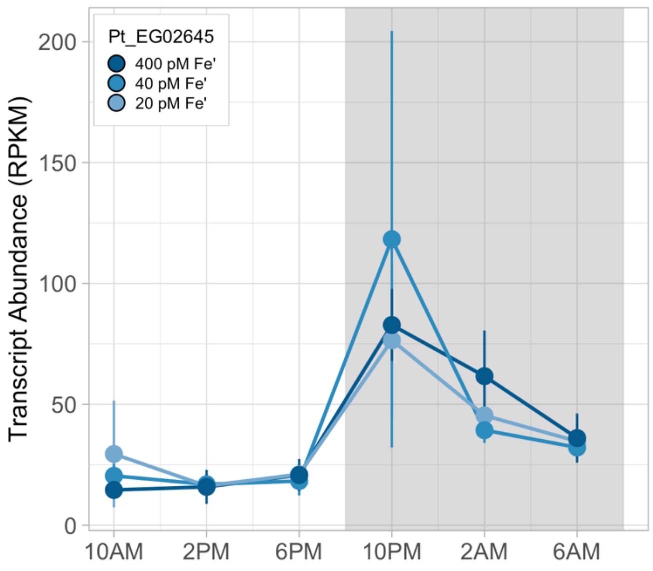

**Supplementary Figure 5. Diel cycle expression dynamics of an orthologous putative antiporter (Phatr3_EG02645) from model pennate diatom *Phaeodactylum tricornutum* (Smith et al., 2016) shows similar to day/night expression as the *T. pseudonana* putative antiporter (262258)**. Averaged transcript abundance (RPKM) is shown over a 24-hr diel cycle at three different iron conditions (20, 40, and 400 pM Fe'). Shaded background indicates the dark period and error bars represent the standard deviation (see Smith et al., 2016 for details).

**Supplementary Table 1.** Primers used in the construction of the final expression vector FCPp-262258-EGFP-FCPt. AttB sites within the primer sequence are indicated in bold.

| Primer name | Sequence (5' to 3') | Gateway site | Direction |
|---|---|---|---|
| 262258_CDS_F | **GGGGACAAGTTTGTACAAAAAAGCAGGCTTA** ATGGCATCCAAACCTGACAACTC | attB1 | Forward |
| 262258_CDS_R | **GGGGACCACTTTGTACAAGAAAGCTGGGTA** AATCCAAACGTACACCACGAGC | attB2 | Reverse |

**Supplementary Table 2.** Top five genes with the largest significant increases and decrease in transcript levels at elevated $CO_2$ (HC vs. LC) from the microarray dataset. Genes with an * had significant differential expression between HC and LC from the RNA-seq dataset (**Supplementary Data File 2**) of the *T. pseudonana* resilience study (Valenzuela et al., 2018).

| Genes with increased expression at HC conditions (i.e., decreased expression at LC) | | | |
|---|---|---|---|
| **Protein Id** | **Annotation** | **Fold Increase** | **FDR** |
| 11637 | PhoD | 22.7 | 3.8E-09 |
| 262258* | putative $Na^{+}(K^{+})/H^{+}$ antiporter | 5.2 | 1.1E-04 |
| 5647* | hypothetical protein | 4.0 | 2.3E-07 |
| 2010 | hypothetical protein | 2.9 | 6.1E-06 |
| 25842 | hypothetical protein | 2.7 | 4.6E-04 |
| Genes with decreased expression at HC conditions (i.e., increased expression at LC) | | | |
| **Protein Id** | **Annotation** | **Fold Decrease** | **FDR** |
| 10917 | hypothetical protein | 6.9 | 2.1E-06 |
| 6528 | zinc transporter (previously reported in a CCM subcluster) | 4.0 | 3.2E-04 |
| 8028 | hypothetical protein | 3.6 | 6.8E-04 |
| 4936 | cathepsin z (Peptidase_C1 superfamily) | 3.4 | 7.0E-04 |
| 10360 | (Tp_bZIP24a) regulator (previously reported in a CCM subcluster) | 3.0 | 3.4E-04 |

**Supplementary Table 3.** *T. pseudonana* putative antiporter (262258) and the *P. tricornutum* ortholog are targeted to the chloroplast (diatom plastid) according to TargetP (v1.1, using 'plant networks') (Emanuelsson et al., 2000).

| ID | Length | cTP | mTP | SP | Other | Loc | RC |
|---|---|---|---|---|---|---|---|
| *T. pseudonana* 262258 | 1067 | 0.867 | 0.005 | 0.060 | 0.490 | C | 4 |
| Phatr3_EG02645 | 967 | 0.919 | 0.024 | 0.006 | 0.484 | C | 3 |

**Description of Supplementary Files:**

**Supplementary Data File 1.** A full list of genes from the three-way ANOVA analysis of transcriptome changes with respect to $CO_2$ level, light level, and growth phase. Differentially expressed genes are split into respective conditions and only gene models with FDR <= 0.001 and a FC > 2 were listed. Genes with multiple ids (e.g., 23654_7354) are the result of potential probes that could in theory be hybridizing against multiple transcripts. There are some cases where this could happen due to very similar or identical 3' regions of genes such as duplicates, paralog genes, or genome assembly issues. This is an artifact of using 3'-directed 'gene-specific' microarrays and were not considered as part of the functional analysis.

**Supplementary Data File 2.** List of significantly differentially expressed genes (n = 101) from the expression analysis of an independent RNA-seq dataset designed to investigate the resilience of *T. pseudonana* at two $CO_2$ levels (300 ppm and 1,000 ppm) over the diel cycle (Valenzuela et al., 2018). See Valenzuela et al., 2018 for detailed expression analysis. The supplementary data file has only the significantly differential expressed genes with a log2 fold-change > 1 when comparing HC (n = 32) vs LC (n = 24) conditions over the day/night cycle during early and late-exponential phase of growth for 3 stages.

**Supplementary Data File 3.** Complete list of FPKMs for all genes (n = 11,780) and conditions (n = 56) from the Valenzuela et al study investigating the resilience of *T. pseudonana* at two $CO_2$ levels (300 ppm and 1,000 ppm) over the diel cycle (Valenzuela et al., 2018).

**Supplementary Data File 4.** Complete list of log2 expression ratios from the microarray dataset for this study (n = 15,059).

**Supplementary Data File 5.** Text file of all antiporter amino acid sequences used in this study for *T. pseuodonana* and *P. tricornutum*. EGFP fusion was performed using the full-length *T. pseudonana* putative antiporter 262258 (1,067 aa), which was PCR amplified from gDNA. Additionally, the *T. pseudonana* full-length antiporter sequence was used for the TargetP analysis. A truncated 262258 gene model (also nested within the full-length sequence) is predicted from the Joint Genome Institute (JGI) *T. pseudonana* assembly (GCA_000149405), which has 439 amino acids, it was used for PredMP transmembrane prediction and Tara Oceans analysis. For the *P. tricornutum* antiporter homolog (Phatr3_EG02645) the gene model from JGI assembly (GCA_000150955.1) predicts 967 amino acids.
