## Supplementary Data File 5 for "A plastid antiporter as a bioindicator of *Thalassiosira pseudonana* resilience"

> *T. pseudonana* putative antiporter 262258 (full-length: 1,067aa)
MASKPDNSSEDVAAVGSEPTNSRARPNRKKVLSFGSPNADFFVIDDNTPASGTRSRQKS
YEGKQRRGSGAASSASPSPINTPELMGPILRRKKVVSMPLGDPRRNDFFYAGDRMGAPY
QANAPVIPMAIDEAASATDNGSEVGHLPGASHFTRYTAGVEATGARARHRERQHTAPA
GRGGFFQIESTGAAMVGTAGGLSGTRGRPNRKKTNSFMAGILARKAVGLDLLPTEEAD
LGVYEFFQPEEVVPEEPPILPEGLDHGEETKEEMEGEKTEEELVEEGESFEEEAVVLKPK
VKLFKPLVFAPKVVPKAGAGGGHGHGGGHGGGGHAPAPSPVQLPEKKTSEKSERTETA
ESSRHSGVVLSASEAVPLTGTTSVSSPQKDADTSEEFEDEESVDLIEKYDDSHIIRATRKD
YFYTAILFAVMSTLVGVVVGWHTRLDESDSIFGPVGLACKTPCRGDIYDQDYFRGQYTF
ATNDVLLLVPHIDTTLTEDSYLKLMIRGVETNQTKWESDDTEFGPASVEGERMSMSVRV
KVDWENPEEPHVIDVISTTGSQQVPPPYYDAYTISGEGAHNDDATHTDDAADHTDDAV
HRVLSSGTTAQQAPDVPHHGELTYNLYAATLHPIAGNSVLIAGLIMIFVYIFILLEVIHRTL
VAIFGSLVALFFFFLMHGGHTESISTLMLHMEWSTLGLLFGMMLLVGELSHTGVFEWCA
VRLLVASKGSFNRLMILLCGLVGLSSAFLDNVTTMLLIAPVTIDMCNILGVDPRPYLIGE
VLLSNVGGTATLIGDPPNIIIGSSFDEVGFVDFIINILPCIFFFCVPASLLLIVKIYKYYLTSE
KMVALDVRQLKKTYQVYDEPRLLIAGTTTFFVILMFFLHPVHHKDTAWIALLGAFITIAF
TNPHDVQDALRNHVEWDTLLFFAGLFVLVEVCAAMGLLQAIGDALASVIKAQPEENQM
TIAITLILWVSAITSAFLDNIPYTATMIPVVKILADELPDTLNIKTLAWALSFGACLGGNGT
LLGASANIVTAGISTSRGYGISFLNFLYPGMLVMIATVAIANIYMLVVYVWIX

> *T. pseudonana* putative antiporter 262258 (JGI gene model: 439aa)
NSVLIAGLIMIFVYIFILLEVIHRTLVAIFGSLVALFFFFLMHGGHTESISTLMLHMEWSTL
GLLFGMMLLVGELSHTGVFEWCAVRLLVASKGSFNRLMILLCGLVGLSSAFLDNVTTM
LLIAPVTIDMCNILGVDPRPYLIGEVLLSNVGGTATLIGDPPNIIIGSSFDEVGFVDFIINILP
CIFFFCVPASLLLIVKIYKYYLTSEKMVALDVRQLKKTYQVYDEPRLLIAGTTTFFVILMF
FLHPVHHKDTAWIALLGAFITIAFTNPHDVQDALRNHVEWDTLLFFAGLFVLVEVCAA
MGLLQAIGDALASVIKAQPEENQMTIAITLILWVSAITSAFLDNIPYTATMIPVVKILADE
LPDTLNIKTLAWALSFGACLGGNGTLLGASANIVTAGISTSRGYGISFLNFLYPGMLVMI
ATVAIANIYMLVVYV

> *P. tricornutum* putative antiporter Phatr3_EG02645 (JGI gene model: 967aa)
MTGDDTTTTPPVVPREAVRPVSSTENAGNYRPRSRQNSRTRQNSRDEMITFNPTTAASM
NNRRVLFSEAARRDGGDNNVAAPEMMGPILRRKKVVSMPLGAGGDFFVSNDVGGPHR
QMLAQYKGVHGDKTTFKGRTKQKSVGAAFVTPADPDPTTVTLPQARIRQDSGTRRDLS
HVIKGVLRRKQVSLDLLGPNDFFALPTDEPEEMVLLPIMPEDDTAQLDDEQGSMTTEKR
LFRPLRFAPRIGVRGNRGGGYGVGHAVEAAPETIPEEAVDEEMVDAFHRYSKTNQDLRL
QVISLKNKLATKSDCSALMEEADSILRTHSELNPDEQLLKRKIMKVQGSIDEEHECSEGID
NNIKDLDNLGIIRATRRDYIKTGILFMIMVALTITVSTWETHLDEESFIFRHVGLACVTEC
RGNLLTRDFFHGHNQFNDGDVIELIMHMDPNSLAETMGALALVQIVGTETNETKAMTT
FGPTAENDRETYDHRLVVNFDRPHEPHIIVVNSTKPNFELSFTLTARLLAPLADNSVAIA
AVIMVVVYLFILLEVIHRTLVAIFGSMVALMFLFVMQNGETESIRQIMLNLEWSTLGLLF
GMMLIVGELSHTGVFEWCAVRLLMASNGSFTRLIVLLCALTAVASAFLDNVTTMLLVA
PVTIDMCNILGVDPRPYLIGEVLLSNIGGTATLIVSLGLVVWVYRYYLTTSTMKVLDTAK
LKTAYPIYDEPRLMIAGTVTAFVIIMFFLHPVHHKDTAWIALLGAFITIAFTNPHDVQDAL
RNHVEWDTLLFFAGLFVLVEACAAMGLLEEIGNLLGDYIQAQEESKQLTLAITLLMWVS

AITSAFLDNIPYTATLIPVIQILADSLPDTLPIEILAWALSFGACLGGNGTLLGASANIVTAG
ISTNKGFEISFLNFLYPGMLFMIVTVAISNLYMLVRYSWI
